# Injury-induced Neuregulin-ErbB signaling from muscle mobilizes stem cells for whole-body regeneration in Acoels

**DOI:** 10.1101/2024.12.23.630141

**Authors:** Brian Stevens, Riley Popp, Heather Valera, Kyle Krueger, Christian P. Petersen

## Abstract

The activation of progenitor cells near wound sites is a common feature of regeneration across species, but the conserved signaling mechanisms responsible for this step in whole-body regeneration are still incompletely understood. The acoel *Hofstenia miamia* undergoes whole-body regeneration using Piwi+ pluripotent adult stem cells (neoblasts) that accumulate at amputation sites early in the regeneration process. The EGFR signaling pathway has broad roles in controlling proliferation, migration, differentiation, and cell survival across metazoans. Using a candidate RNAi screening approach, we identify the *Hofstenia* EGFR *erbB4-2* and Neuregulin *nrg-1* genes as essential for blastema formation. Structure prediction of NRG-1 and ERBB4-2 proteins supports the likelihood of these factors interacting directly. After amputation injuries, *nrg-1* expression is induced in body-wall muscle cells at the wound site by 6 hours and localizes to the tip of the outgrowing blastema over the next several days, while *erbB4-2* is broadly expressed, including in muscle and neoblasts. Under *nrg-1(RNAi)* and *erbB4-2(RNAi)* conditions that impair blastema formation, animals still undergo the earliest responses to injury to activate expression of the Early Growth Response transcription factor *egr*, indicating a crucial role for EGFR signaling downstream of initial wound activation. *nrg-1(RNAi)* and *erbB4-2(RNAi)* animals possess Piwi+ and H3P+ mitotic neoblasts which hyperproliferate normally after amputation, but these cells fail to accumulate at the wound site. Therefore, muscle provides a source for Neuregulin-ErbB signaling necessary for the mobilization of proliferative progenitors to enable blastema outgrowth for whole-body regeneration in *Hofstenia*. These results indicate a shared functional requirement for muscle signaling to enable regeneration between planarians and acoels across 550 million years of evolution.

## Introduction

Many regenerative organisms respond to injury by mounting a rapidly induced transcriptional program that promotes blastema outgrowth and tissue repair (Gehrke et al., 2019; Wenemoser et al., 2012; Wurtzel et al., 2015). A common hallmark of this process involves the proliferative activation of stem cells and other latent progenitors, or activation of proliferative progenitors through dedifferentiation, and their targeting to wound sites (Poss and Tanaka, 2024). Injury-induced proliferation occurs across a wide variety of regenerative organisms (Ricci and Srivastava, 2018), including axolotls (Johnson et al., 2018), zebrafish (Poss et al., 2000), planarians (Newmark and Sanchez Alvarado, 2000; Wenemoser and Reddien, 2010), acoels (Srivastava et al., 2014), and Cnidarians (Amiel and Rottinger, 2021; Bradshaw et al., 2015). The activation of progenitors is critical for regeneration competence, and similarity in these processes across regenerative organisms could point to a set of common mechanisms to promote regeneration. The detailed mechanisms relaying wound information to the effector steps of blastema formation in whole-body regeneration are still incompletely understood.

The acoel *Hofstenia miamia* is an emerging model organism to study the mechanistic basis for whole-body regeneration (Srivastava, 2022). This species undergoes whole-body regeneration through activation of conserved injury-induced responses (Gehrke et al., 2019), the use of Piwi+ neoblast pluripotent stem cells to form all adult tissue types (Kimura et al., 2022; Srivastava et al., 2014), and control of body patterning through conserved Wnt/BMP signals (Srivastava et al., 2014) produced from body-wall muscle (Raz et al., 2017). The availability of transgenesis in this organism (Ricci and Srivastava, 2021), a comprehensive single-cell transcriptome atlas (Hulett et al., 2023), systemic RNAi (Srivastava et al., 2014), and detailed information about embryogenesis (Kimura et al., 2022; Kimura et al., 2021), organ systems (Hulett et al., 2020), and differentiation pathways (Hulett et al., 2024) make *Hofstenia* a powerful system to understand whole-body regeneration mechanisms. In addition, the unique phylogenetic placement of acoels as either basal Deuterostomes (Philippe et al., 2011) or basal Bilaterians (Cannon et al., 2016; Hejnol et al., 2009; Ruiz-Trillo et al., 1999; Srivastava et al., 2014) has enabled the use of this organism for comparative approaches to uncover deeply conserved mechanisms of regeneration (Gehrke et al., 2019; Poulet et al., 2023; Raz et al., 2017; Srivastava et al., 2014; Tewari et al., 2019).

*Hofstenia* initiates regeneration through rapidly inducing the expression of the Early Growth Response transcription factor *egr* by 1 hour post-amputation, which acts as a pioneer factor to activate expression of several downstream genes and is essential for regeneration. ATACseq and RNAi established that EGR protein likely directly activates the expression of several downstream genes activated by 6 hours, including *runt*, *follistatin-like (fstl), nrg-1, nrg-2, deaf1, nlk, ankrd, mtss-1, p-protein,* and *wie-1* (Gehrke et al., 2019). Several components of this pathway have conserved involvement in promoting whole-body regeneration. For example, Runt transcription factors are activated by injury and required for regeneration in planarians *Schmidtea mediterranea* and acoels *Hofstenia* and *Convolutriloba longfissura* (Gehrke et al., 2019; Nanes Sarfati et al., 2024; Wenemoser et al., 2012), and *follistatin* family members are also required for regeneration in planarians (Gavino et al., 2013; Roberts-Galbraith and Newmark, 2013). In *Hofstenia,* single-cell RNAseq found that several key injury-induced are expressed strongly from muscle (including *egr, runt, nrg-1, fstl*), and also neurons and/or neoblasts (*p-protein, nlk1, deaf1)* and epidermal progenitors (*nrg-2*) (Hulett et al., 2023). *egr* and *runt* are functionally required for blastema formation (Gehrke et al., 2019). However, the functional roles of other downstream genes to promote specific regeneration activities and pathways have not yet been determined. In addition, the functional role of muscle in the early injury response has not yet been established.

EGF (Epidermal Growth Factor) signaling has diverse roles in regeneration across species. Members of the ErbB family of EGF receptors (EGFRs), as well as their associated EGF family ligands, including EGF and neuregulins (NRGs), have been demonstrated to play a functional role in promoting cell proliferation and blastema formation in a variety of regenerative vertebrate systems, including in zebrafish heart (Gemberling et al., 2015), zebrafish fin (Rojas-Munoz et al., 2009), and axolotl limb regeneration (Farkas et al., 2016). In planarians, a neuregulin and EGFR pair control asymmetric division and clonal growth of neoblasts repopulating the animal after sublethal irradiation (Lei et al., 2016), and EGFRs are required for regeneration and homeostasis of several organ systems (Barberan and Cebria, 2019; Barberan et al., 2016; Emili et al., 2019; Fraguas et al., 2011; Rink et al., 2011).

Based on the widespread use of EGF signaling, we used a candidate approach to comprehensively examine EGFR signaling pathway ligands and receptors in *Hofstenia* and determine their functional roles in regeneration. We identify a causative role for *nrg-1* and *erbB4-2* in controlling the injury-induced mobilization of progenitor cells necessary for subsequent blastema formation. *nrg-1* is expressed exclusively from body-wall muscle at the injury site and also within the distal blastema over the course of several days, indicating muscle is a source of EGF signaling necessary to enable outgrowth.

## Results

To examine candidate functions for EGFR signaling in *Hofstenia*, we cloned pathway receptors and ligands and performed candidate screening by RNAi. Blast searching of the *Hofstenia* transcriptome revealed matches to two possible EGF receptors annotated as *erbB4* (referred here as *erbB4-1*) and *erbB4-2* (Fig. S1A). Both genes encoded an intracellular tyrosine kinase domain, but differed in the domain structure of their extracellular regions. *erbB4-2’s* ectodomain encoded a canonical set of Receptor L and Furin domains typical of canonical EGFRs (Ferguson, 2008). By contrast, *erbB4-1’s* extracellular region was truncated and lacked Receptor L domains, and contained an unannotated region that Foldseek predicted to have structural similarity to a divergent CUB domain, which are found in a variety of extracellular proteins outside of the canonical EGF pathway (Bork and Beckmann, 1993). Therefore, *erbB4-1* is unlikely to interact directly with canonical EGFR family ligands. We tested for roles of these genes in *Hofstenia* regeneration using RNA interference by microinjection of dsRNA into body cavities on three sequential days, followed by transverse amputations. Both the head and tail fragments from *erbB4-2(RNAi)* animals completely failed to form regeneration blastemas by day 7 (57/62 head fragments, 56/58 tail fragments), whereas animals treated with control dsRNA all succeeded at regeneration (Fig. 1A). Therefore, *erbB4-2* is essential for regeneration in *Hofstenia*. By contrast, the majority of animals fragments treated with dsRNA targeting *erbB4-1* through this schedule succeeded at regeneration (9/11 head fragments and 10/12 tail fragments) (Fig. 1A). Because of the divergent ectodomain and lack of ascribed function in regeneration, we did not consider further roles for *erbb4-1*.

**Figure 1.**
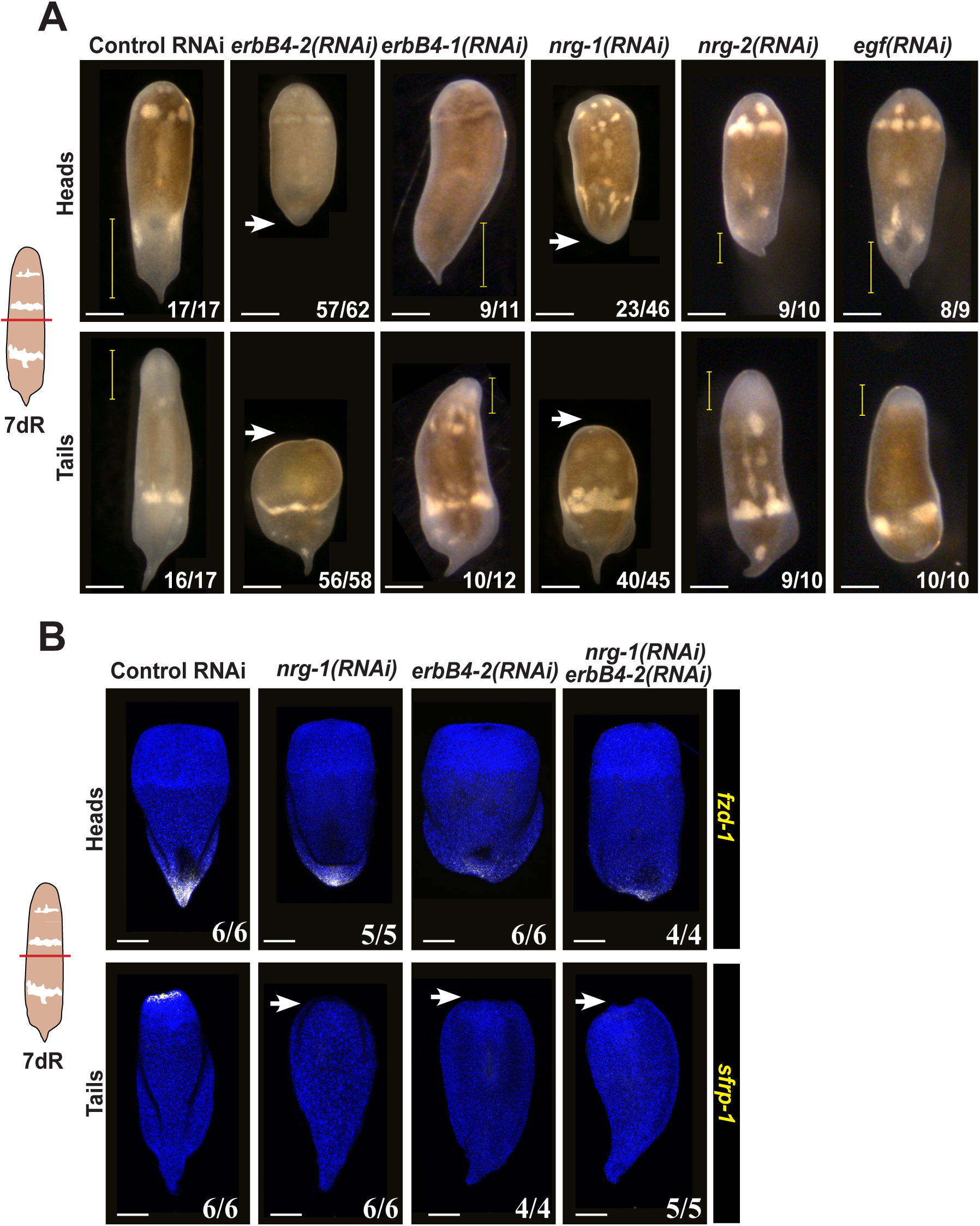
*Hofstenia erbB4-2* and *nrg-1* are required for regeneration. (A) Animals were injected with the indicated dsRNAs for three sequential days gene prior to transverse amputations to examine regeneration of head and tail fragments. Control animals, *erbB4-1(RNAi)*, and *nrg2(RNAi)* and *egf(RNAi)* animals formed normal regeneration blastemas (yellow brackets), whereas *erbB4-2(RNAi)* and *nrg-1(RNAi)* animals lacked a blastema (white arrows). (B) FISH to detect expression of posterior marker *fzd-1* in head fragments and *sfrp-1* in tail fragments. Single or double RNAi of *erbB4-2* and *nrg-1* prevented new expression of *sfrp-1* in tail fragments (arrows) and weakened but still permitted *fzd-1* expression in head fragments under these conditions. Numbers indicate scoring of animals representing the outcome found in each representative image across several experiments. Scale bars, 500 (A) or 200 microns (B).

We next sought EGF family ligands which might signal through *erbB4-2* to promote regeneration. *Hofstenia* encodes two previously annotated Neuregulin family homologs, *nrg-1* and *nrg-2* (Gehrke et al., 2019), and one EGF homolog *egf* (Fig. S1B). EGF family members undergo proteolytic maturation of the extracellular EGF domain-containing regions to enable formation of signaling competent secreted ligands. NRG-1 has a domain structure similar to *Drosophila* Vein family members, possessing only a single EGF domain along with an adjacent transmembrane domain. NRG-2 possesses an IG and EGF ectodomain similar to the ectodomain of other Neuregulin factors such as human NRG1beta, and the domain structure of the protein encoded by Hofstenia *egf* resembles known EGF protein families. Any or all of the three *Hofstenia* EGFs could potentially mediate the pro-regenerative effects of *erbB4-2*, and we tested this hypothesis using RNAi to try to phenocopy the effects of *erbB4-2* inhibition. Using the same schedule for RNAi described above, *nrg-1* inhibition completely prevented blastema formation at a high penetrance (40/45 tail fragments and 23/46 head fragments failed to regenerate).

Conversely, *nrg-2(RNAi)* and *egf(RNAi)* animals all regenerated normally (Fig. 1A). Therefore, the similar regeneration deficient phenotypes after *nrg-1* and *erbb4-2* RNAi suggest these factors likely participate in a common pathway to promote regeneration. To examine the character of regeneration deficiency under these treatments, we stained fragments for head and tail markers in order to assess the regeneration defect histologically (Fig 1B). Tail fragments from single or double RNAi of *nrg-1* and *erbB4-2* completely failed to express anterior marker *sfrp-1*, whose re-expression in *Hofstenia* regeneration is known to require neoblast activity (Raz et al., 2017). Despite failing to grow blastema tissue, head fragments under these conditions re-expressed low levels of *frizzled-1,* a posterior marker known to be capable of re-activation in regeneration from pre-existing muscle cells independent of stem cells (Raz et al., 2017) downstream of *egr* and *wnt-3* (Ramirez et al., 2020). Therefore, *nrg-1* and *erbB4-2* RNAi conditions caused a regeneration deficient phenotype similar to loss of neoblasts.

To further examine the plausibility of NRG-1/ERBB4-2 protein interactions, we used Alphafold3 to predict the structures of NRG-1 in isolation and also in complex with the ERBB4-2 ectodomain. The EGF domain from *Hofstenia* NRG-1 protein sequence aligned with several Neuregulin homologs showing conservation of characteristic cysteine residues (Fig S2A). In addition, Alphafold3 predicted a similar overall fold between *Hofstenia* NRG-1 and human NRG-1beta, with majority of the sequence having high confidence (>70 pIDDT) (Fig S2B).

Alphafold3 also predicted an NRG-1 binding pocket within ERBB4-2’s canonical ectodomains I and III (Fig S2C-E), at a position analogous to known Neuregulin/ERBB4 ligand/receptor binding interactions (Ferguson, 2008). Together, this computational structure prediction supports a model in which NRG-1 signals directly through ERBB4-2 to promote regeneration.

RNAseq and FISH previously identified *nrg-1* expression as injury induced by 6 hours post-amputation (Gehrke et al., 2019). However, the detailed expression of this gene throughout regeneration has not been reported. Using FISH, we detected expression of *nrg-1* in dispersed cells at 6 hours near the wound site, followed by stronger expression localized to the distal tip of the blastema for the next several days and lowering at 5 days (Fig 2A-B). The expression of *nrg-1* both at wound sites and later within the regenerating blastema suggests it is responsible for an active signaling process required during the process of outgrowth.

**Figure 2.**
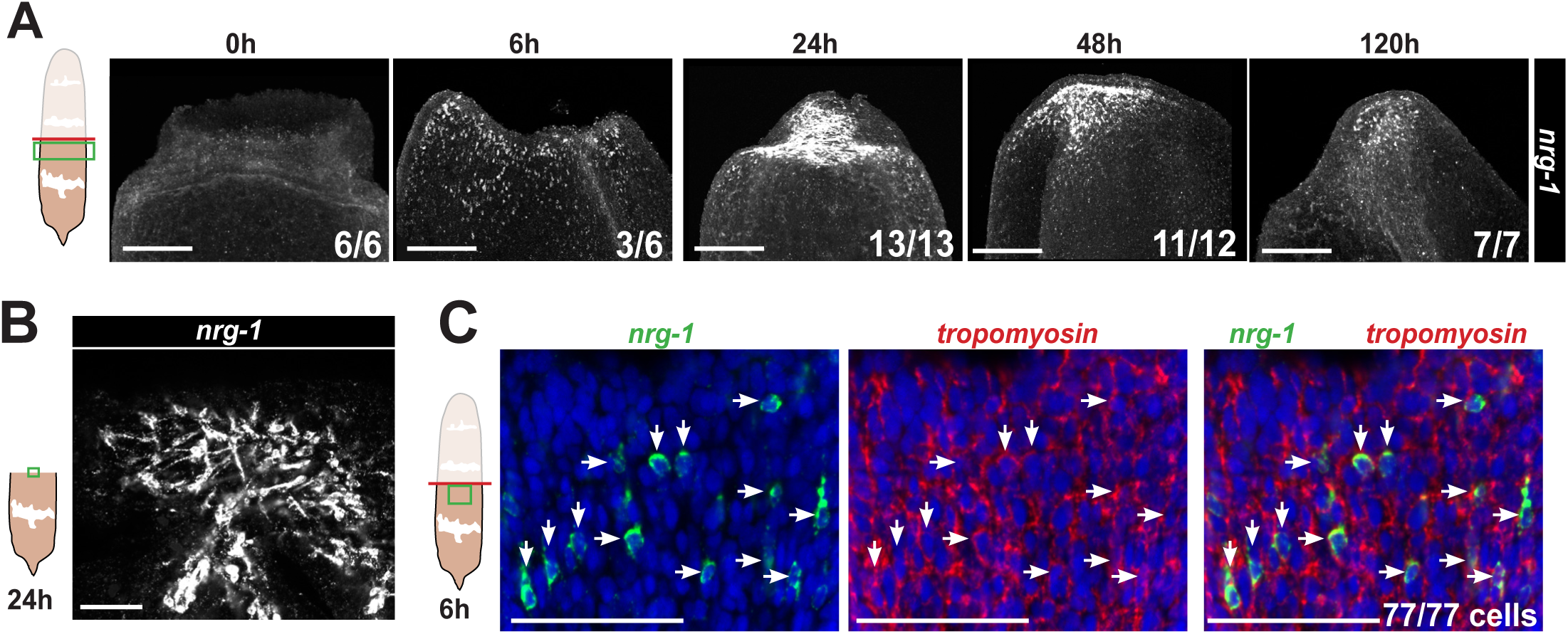
Wound-induced *nrg-1* expression marks muscle and the terminus of the outgrowing blastema. Cartoons indicate surgeries (red lines) and regions imaged (green boxes). (A) FISH detecting *nrg-1* mRNA in regenerating *Hofstenia*. *nrg-1* was expressed in dispersed muscle at 6-hours post amputation. By 24 hours, *nrg-1* expression was present more strongly and distributed throughout the blastema, then coalesced to the distal region of the blastema by 48 hours, followed by a reduction by 120 hours. Images show maximum projections from confocal z-stacks. (B) Single confocal z-slice of *nrg-1* expressing cells in the blastema from a 24-hour regenerating animal, acquired at 20x magnification with confocal zoom. (C) Double FISH detecting expression of *nrg-1* in *tropomyosin*+ muscle cells at 6 hours (77/77 *nrg1+* cells). Image shows a confocal z-slice in the plane of body-wall muscle soma acquired at 20x magnification and 5.46x confocal zoom. Scale bars = 200 (A) and 50 (B, C) microns.

To gain insights into the signal sending and receiving cells relevant for this regulation, we examined the expression of *nrg-1* and *erbB4-2*. Prior single-cell RNAseq identified *nrg-1* as strongly enriched in muscle from uninjured animals and also in muscle at 6 hours post amputation (Hulett et al., 2023). Mining this cell atlas, we found that *erbB4-2* expression was broad, including in both muscle and neoblasts (Fig S3). Using FISH, we were not able to detect any signal for *erbB4-2* transcript compared to background, perhaps due to its broad and weak expression levels. However, double-FISH verified that *nrg-1* is indeed expressed from *tropomyosin+* muscle at wound sites (Fig 2C). We counted *nrg-1+* cells in whole-animal z-stacks and found that 100% co-expressed *tropomyosin* at 6 hours (77/77 cells), indicating muscle is the sole source for the activation of this gene after injury. Consistent with these findings, *nrg-1-*expressing cells of the distal blastema appeared as a reticulated mesh with projections reminiscent of muscle fibers (Fig 2B). Therefore, *nrg-1* constitutes a wound-induced signal produced exclusively from muscle to drive subsequent blastema formation.

We performed experiments to determine the step(s) of regeneration affected by inhibition of *nrg-1* and *erbB4-2* under these conditions. Injuries initially activate expression of the *egr* transcription factor by 1 hour post-amputation, which subsequently binds and activates numerous wound-induced target genes, including *nrg-1* by 6 hours. Consistent with this model, using the same dsRNA dosing schedule that resulted in a highly penetrant phenotype of defective blastema formation (Fig 1A), tail fragments undergoing single and double RNAi of *nrg-1* and *erbB4-2* still expressed high levels of the transcription factor *egr* at the wound site at 6 hours (Fig 3A). By contrast, *egr* expression at the wound site was undetectable in *egr(RNAi)* animals. In regenerating head fragments undergoing *nrg-1* or *erbB4-2* RNAi, the expression of *egr* was strongly detected but in some samples appeared somewhat less abundant, particularly in *nrg-1;erbB4-2*(RNAi) animals. These results could point to the capability for *nrg-1* to engage in autoregulation in some contexts. However, the data from tail fragments indicate that *nrg-1* and *erbB4-2* execute a critical pro-regenerative role downstream of *egr* expression activation.

**Figure 3.**
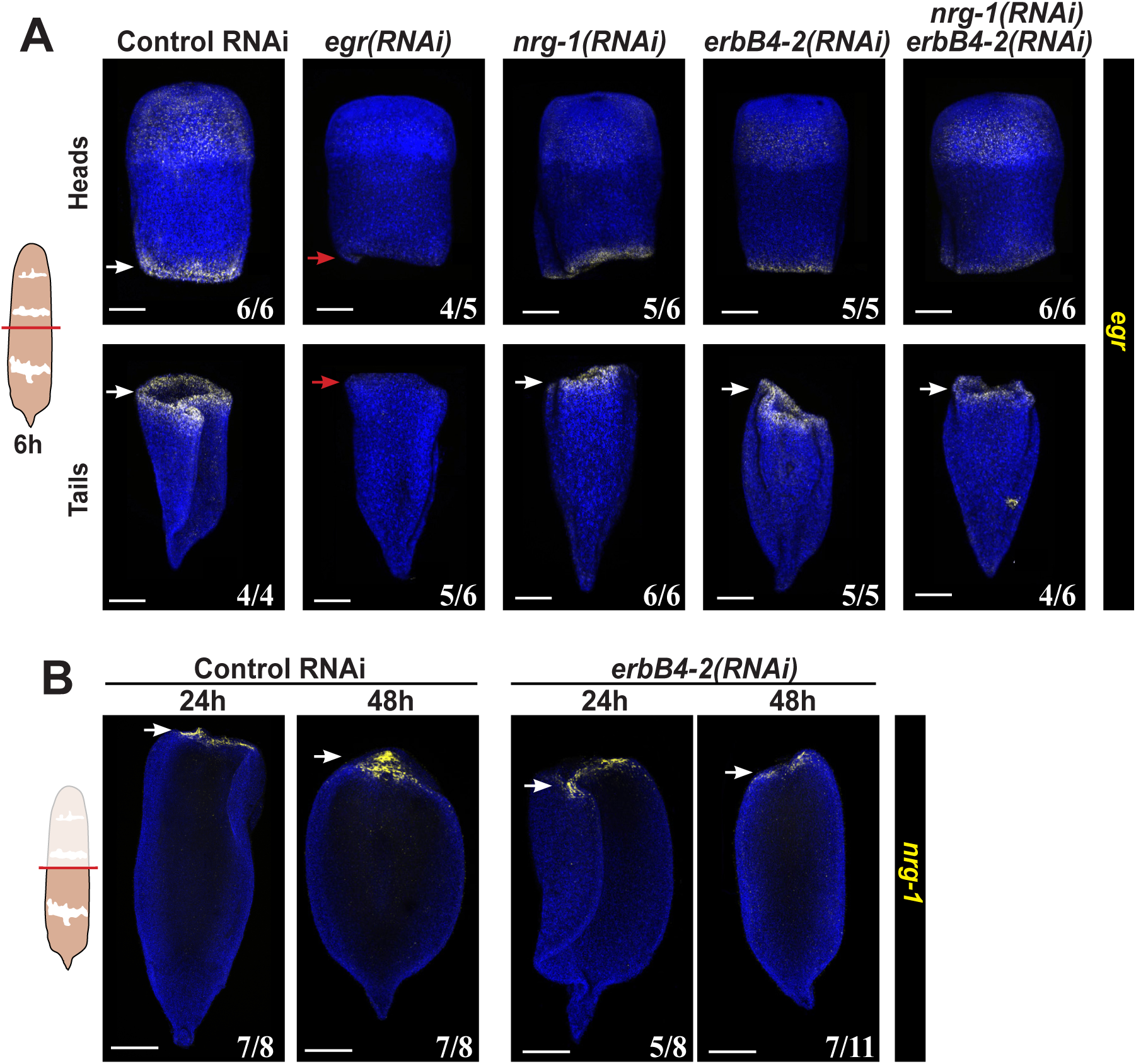
*nrg-1* and *erbB4-2* have an essential role in regeneration downstream of injury-induced *egr* activation. (A) FISH to detect expression of *egr* following the indicated RNAi treatments. *egr* expression was detected at the wound site in control animals (white arrows), but was absent from *egr(RNAi)* animals (red arrows). Under these conditions, *nrg-1(RNAi)* and *erbB4-2(RNAi)* treatments enabled normal levels of *egr* expression in tail fragments (white arrows). Substantial wound-proximal *egr* expression was still detected, though at somewhat reduced intensity, in regenerating head fragments. (B) *erbB4-2(RNAi)* animals expressed wound-induced *nrg-1* for several days. Scale bars, 200 microns. Scorings indicate the number of animals representing the images shown.

EGR transcription factors themselves can act downstream of EGFR signal cascades (Citri and Yarden, 2006), so we further tested whether *erbB4-2* might be required to sustain *nrg-1* expression in the blastema through a positive feedback loop. However, under *erbB4-2(RNAi)* conditions that cause fully penetrant regeneration deficiency, *nrg-1* expression nonetheless continued at the wound site for several days (Fig. 3B). We cannot rule out the possible involvement of long-perduring populations of ERBB4-2 protein that could contribute to regeneration in some way. However, our identification of conditions in which *nrg-1* and *erbB4-2* knockdown strongly impairs regeneration without affecting early injury-induced expression of *egr* and *nrg-1,* respectively, argue that these signals mediate an essential step of regeneration downstream of the process of initial wound activation.

Neuregulin and ErbB can act as mitogens across species (Citri and Yarden, 2006), suggesting the possibility that *nrg-1* and *erbB4-2* might promote the proliferation of neoblasts in *Hofstenia*. *nrg-1* and *erbB4-2* RNAi did not eliminate *piwi-1+* neoblasts, suggesting a more specific role in the animal, at least under these conditions (Fig. 4A). However, regenerating tail fragments under these treatments failed to accumulate *piwi-1+* cells at the wound site (Fig. 4A). Using anti-H3P staining, we quantified effects of *nrg-1* and *erbB4-2* RNAi on mitotic cells in regenerating tail fragments, which undergo a re-localization to the wound site by 72-hours during the process of blastema outgrowth (Fig. 4B-D). Overall numbers of mitotic cells were not altered by *nrg-1* or *erbB4-2* RNAi at 0 hours and 6 hours, and the global increase of proliferative cell numbers by 72 hours was not affected (Fig. 4B). However, *nrg-1* and *erbB4-2* inhibition completely prevented the localized accumulation of H3P+ cells near the wound site by 72 hours (Fig. 4B, D). Therefore, we did not detect evidence for *nrg-1* and *erbB4-2* to control proliferation in general, but find they are critical for the localization of proliferative cells at the onset of the regeneration process.

**Figure 4.**
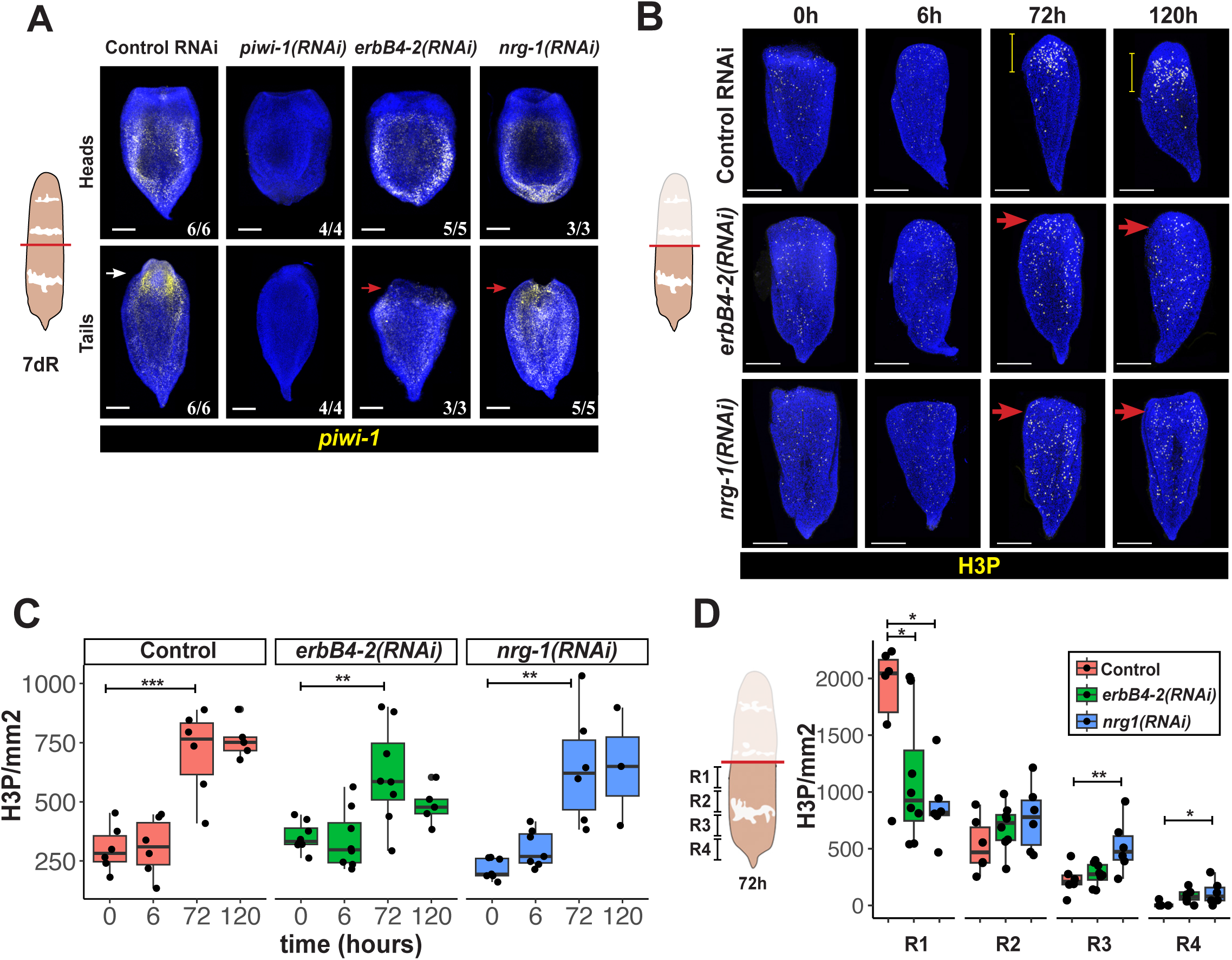
*nrg-1* and *erbB4-2* are required to mount a localized proliferative response during blastema formation. (A) FISH of the neoblast marker *piwi-1* in regenerating head and tail fragments after indicated RNAi treatments. *piwi-1* RNAi eliminated the staining from *piwi-1* riboprobe. *piwi-1* expression persisted after *nrg-1* and *erbB4-2* RNAi, but failed to localize to the blastema in regenerating tail fragments (red arrows) compared to control animals (white arrow). (B) Detection of mitotic cells in regenerating tail fragments by immunostaining with anti-H3P antibody. In *nrg-1* and *erbb4-2(RNAi),* mitotic distributions appeared normal at 0 and 6 hours but failed to localize to the wound site by 72 hours (red arrows) compared to control animals which successfully localized mitotic cells (yellow bars). (C-D) Quantification of mitotic cells from (B) normalized to animal area, with individual datapoints showing enumerations for separate experimental samples. (C) *nrg-1* and *erbB4-2* RNAi still enabled a global elevation of mitotic cell numbers at 72 hours, similar to control animals (*** p<0.001, ** p< 0.01, Kruskal-Wallis test followed by Dunnett’s test). Additionally, mitotic cell numbers were not significantly different across the populations at 0h or 72h (p=0.32 for *erbB4-2(RNAi)* versus control at 0h, p=0.07 for *nrg-1(RNAi)* versus control at 0h, p=0.63 for *erbB4-2(RNAi)* versus control at 72h, p=0.83 for *nrg-1(RNAi)* versus control at 72h by Kruskal-Wallis test and Dunnett’s test to compare means). (D) Analysis of the spatial distribution of mitotic cells across evenly spaced A/P regions (R1, R2, R3, R4) confirmed the defective wound proximal localization of H3P+ cells in *nrg-1* and *erbB4-2(RNAi)* animals at 72 hours (** p<0.01, * p<0.05, by Kruskal-Wallis test and Dunnett’s test to compare means across multiple populations). Scale bars, 200 microns.

## Discussion

These experiments suggest a model for participation of *nrg-1* signaling through *erbB4-2* in the process of blastema outgrowth necessary for whole-body regeneration in *Hofstenia* (Fig. 5). An upstream injury signaling process activates *nrg-1* in muscle near the wound site at early times (6 hours) and later becomes expressed within blastema to increasingly distal regions (1-5 days). *nrg-1* signals through *erbB4-2* to localize proliferating progenitor cells and initiate blastema differentiation and formation. *erbB4-2* is expressed broadly, including in neoblasts and muscle. Therefore, NRG-1 could either signal to neoblasts directly, or perhaps through an indirect mechanism involving signal reception in another cell type, for example muscle itself, which subsequently affects neoblast targeting and proliferation needed for blastema formation (Fig. 5A).

**Figure 5.**
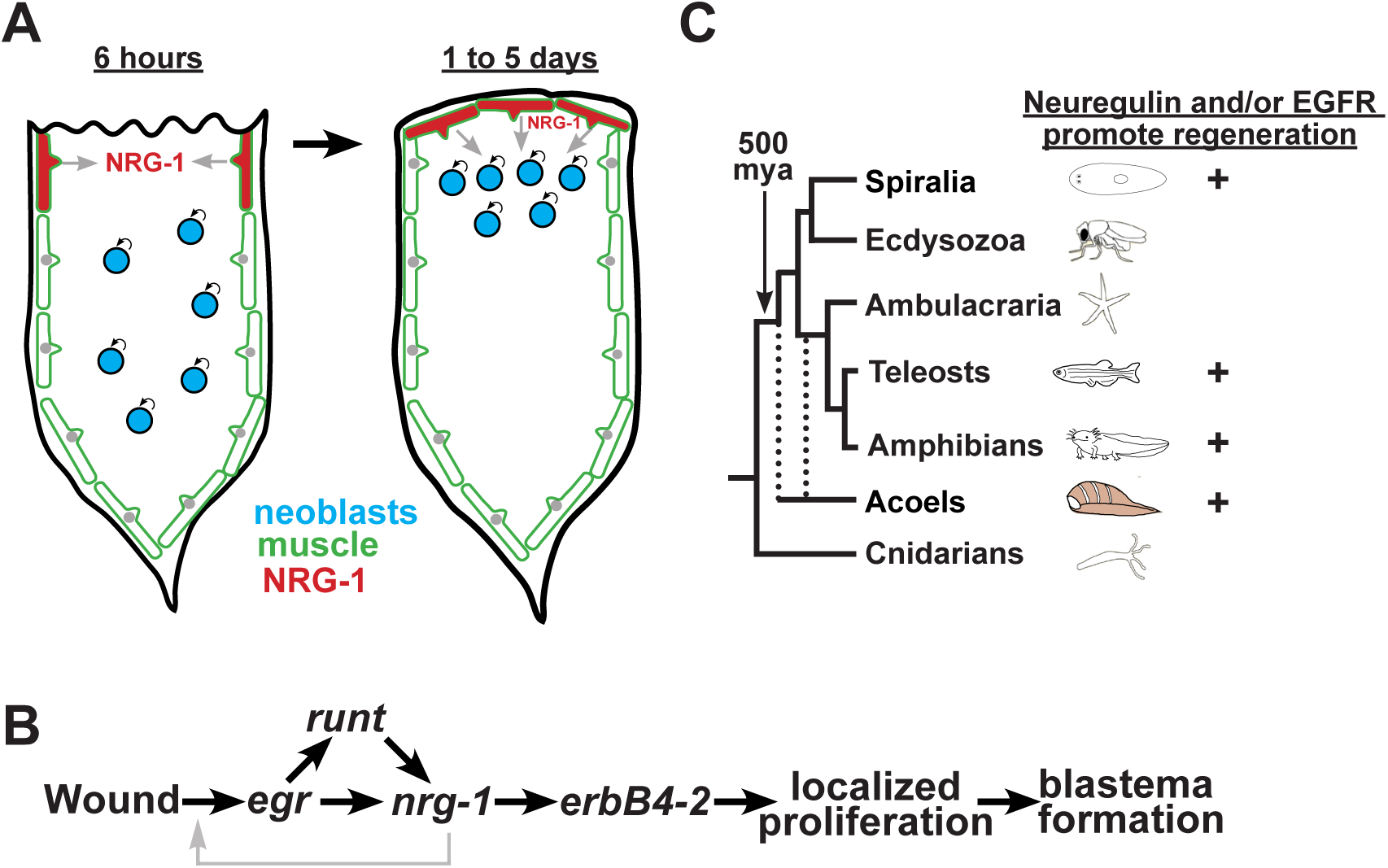
Model. (A) Early wound signaling activates expression of *nrg-1* in muscle near the wound site by 6 hours and subsequently in the distal blastema region from 1-5 days. As regeneration progresses, *nrg-1* expressed from muscle signals through *erbB4-2*, and receiving cells of this signaling could either be neoblasts (gray arrows), muscle, or other cell types. A downstream consequence of the signaling results in localization of neoblast proliferation near the blastema region, which correlates in space and time approximately with the locations of *nrg-1*’s late expression phase. The activity of *nrg-1* and *erbB4-2* promotes blastema outgrowth. (B) Regulatory model identifying a critical role for *nrg-1* and *erbB4-2* functionally downstream of *egr*. Wounding activates immediate early expression of *egr,* which is responsible for transcriptional activation of *nrg-1* and *runt-1. nrg-1* is functionally required for regeneration and acts through *erbB4-2,* and these factors exert critical responses downstream of *egr* in order to localize proliferation to wound sites and also cause blastema outgrowth. Gray lines: contributions from *nrg-1* signal feedback and/or roles for pre-existing NRG-1 protein in the wound activation process (see text). (C) Phylogeny showing the conserved use of Neuregulin and/or EGFR signaling to functionally promote regeneration. Acoels are believed to be either early-branching Deuterostomes within Xenacoelomorpha or form a sister group to all Bilaterian animals (dotted lines).

Several lines of evidence support a critical step of *nrg-1* action downstream of early wound activation (Fig. 5B). First, prior work showed that *egr* and *runt-1* are necessary for the activation of *nrg-1* expression following injury (Gehrke et al., 2019). Second, *nrg-1* expression in regeneration (by 6 hours) occurs somewhat later than *egr* expression (by 1 hours) (Gehrke et al., 2019). Additionally, in examining *nrg-1* expression throughout regeneration, this factor has ongoing expression in the blastema for several days, suggestive of a likely function for new expression of *nrg-1* in regeneration (Fig. 2). We used a relatively short RNAi dosing strategy to assess the role of *nrg-1* in regeneration (injections of dsRNA on 3 consecutive days prior to amputation). Under these conditions, *nrg-1* silencing blocked regeneration to a strong degree in regenerating tail fragments (Fig. 1A-B) but enabled expression of *egr* at 6 hours (Fig. 3A), thus demonstrating a critical role for this gene downstream of *egr* activation and linked to the subsequent process of blastema outgrowth. Although blastema outgrowth completely failed in tail fragments undergoing *nrg-1* RNAi and these fragments did not produce any new *sfrp1+* cells, head fragments still succeeded at some re-expression of *fzd-1* (Fig. 1B). *fzd-1* re-expression can still occur in irradiated animals lacking stem cells and represents the output of early polarity decisions mediated by injury-induced Wnt signaling in pre-existing muscle (Raz et al., 2017).

Notably, the process of *fzd-1* reactivation at posterior-facing wounds requires *egr* to activate wound-induced expression of the polarity factor *wnt-3* (Ramirez et al., 2020), indicating that in our experiments, *nrg-1* silencing still permitted an *egr-*dependent process. Together, these experiments demonstrate that new expression of *nrg-1* regulates a critical step of regenerative outgrowth.

A recent study reported an additional critical role for NRG-1-dependent phospho-ERK activation early after wounding (Breen and Srivastava, 2024). Phospho-ERK (pERK) activates at a very early stage following injury (starting prior to 10 minutes and ending by 12 hours), and inhibition of ERK signaling by treatment with the U0126 pharmacological inhibitor of MEK prevented injury-induced new expression of *egr, nrg-1*, and *runt*, and also delayed overall regeneration progress (Breen and Srivastava, 2024). Neuregulins in other systems can signal through pERK, so the study also examined whether long-term knockdown of *nrg-1* could deplete any pre-existing NRG-1 protein that might be involved in the activation of early pERK activation upstream of subsequent *egr* transcriptional activation. Using a longer RNAi schedule to inhibit *nrg-1* dsRNA homeostatically (over 12 days), amputated animals were deficient at early phospho-Erk activation, supporting a likely role for pre-existing NRG1 to become activated by injury (Breen and Srivastava, 2024). In our experiments reported here, a shorter dose schedule of *nrg-1* dsRNA (over 3 days) still enabled wound-induced expression of *egr* but fully prevented regeneration in tail fragments. Given that *egr* activation occurs downstream of pErk (Breen and Srivastava, 2024), it is likely our shorter RNAi dosing schedule would have relatively less influence on pre-existing NRG1 protein. In addition, new expression of *nrg-1* after injury depends on *egr* (Gehrke et al., 2019), and we find that *nrg-1* expression is sustained in the blastema of normal animals for several days after the end of the initial 12-hour wave of pErk activation following injury (Breen and Srivastava, 2024). Together, the two studies reveal complementary information showing the participation of NRG1 in mediating both the very earliest responses to wounding (denoted by the gray arrows in Fig. 5B) and also driving an essential process downstream of early wound activation.

The sustained expression of *nrg-1* at the distal tip of the blastema near the accumulation of mitotic neoblasts over several days is suggestive of an ongoing role this factor plays subsequent to the early wound activation process. This expression phase is also correlated with the location and timing of neoblast localization to the wound site (by 48-72 hours). The process of progenitor accumulation could occur through either local activation of proliferation, or perhaps through migratory targeting of proliferative cells. In our experiments, *nrg-1* and *erbB4-2* RNAi did not cause a failure to globally increase the number of mitotic cells after injury (total mitotic cells at 0 hours versus 72 hours) (Fig. 4C), suggesting these factors are likely not responsible for all cell proliferation. The normal process of progenitor accumulation also depletes H3P+ cells further away from the wound site (Fig. 4B). Therefore, if *nrg-1* and *erbB4-2* do act to promote local proliferation near the blastema, an additional process would have to account for the fact that H3P+ cells away from the wound site remain present after the inhibition of these factors. Instead, if *nrg-1* drives neoblast accumulation near the blastema through regulating migration, this model could account for the lack of changes to overall mitotic numbers and failure for them to gather in the correct locations. Neuregulin/ErbB4 signaling can direct proliferation (Lemke and Brockes, 1984), differentiation (Plowman et al., 1993), or cell migration (Rio et al., 1997), and the accumulation of proliferative cells at site of regeneration can involve both proliferation (Newmark and Sanchez Alvarado, 2000) and/or migration (Tahara et al., 2016). Future work will be necessary to fully resolve these or other possible models for how *nrg-1* signals downstream to influence progenitor behavior and initiates blastema formation in *Hofstenia*.

Our study also furthers prior work showing the importance of muscle as a source of signals needed for regeneration. In planarians, body-wall muscle expresses Wnt, BMP, and Hox gene encompassing a coordinate system that patters the body axis in three dimensions (Witchley et al., 2013). Injured muscle expresses pro-regenerative factors *follistatin* and *notum* (Scimone et al., 2017), ablation of specific muscle subsets through *myoD*, *nkx1-1,* and *foxF-1* RNAi prevents or mispatterns regeneration (Scimone et al., 2017; Scimone et al., 2018), and disruption of muscle transcription factors such as *alx-3* can also prevent formation of signaling centers needed for regeneration in this species (Akheralie et al., 2023). In addition, planarian muscle is required for a rapid and body-wide activation of phospho-Erk, suggesting that communication systems across these elongated cells mount the systemic response driving whole-body regeneration (Fan et al., 2023). Planarian muscle also provides the majority of extracellular matrix necessary for body integrity (Cote et al., 2019), and express factors such as *collagen IV* (Chan et al., 2021) and *hemicentin* (Cote et al., 2019; Lindsay-Mosher et al., 2020) that can support or constrain stem cell function. In *Hofstenia,* muscle is a primary source of body-wide positional information (Raz et al., 2017) and expresses several conserved injury-induced genes (Hulett et al., 2023). Given the high sequence divergence between planarians and acoels, which may likely diverged over 550 million years ago, these observations suggest a potentially ancient use of muscle for executing key functions in regeneration within Bilaterians. Our analysis, along with a complementary study (Breen, 2024), demonstrate that a *Hofstenia* muscle functionally enables blastema outgrowth by expression and activity of *nrg-1.* In addition, the biphasic pattern of *nrg-1*’s expression activation program near the wound site and then later within the blastema is also reminiscent of the patterns of activation of muscle-expressed signaling factors that promote regeneration in planarians, for example *wnt1* (Petersen and Reddien, 2009), *notum* (Petersen and Reddien, 2011), and *follistatin* (Gavino et al., 2013; Roberts-Galbraith and Newmark, 2013).

These factors all activate within pre-existing muscle cells at early times after amputation, followed by a sustained activation in newly differentiated muscle cells of the blastema (Scimone et al., 2014; Vasquez-Doorman and Petersen, 2014; Vogg et al., 2014). The commonality of biphasic wound-induced expression from muscle could indicate common processes involved in sustaining pro-regenerative signaling for tissue assembly across the timescales needed for growth.

Together, our results identify an exclusively muscle-expressed, injury-induced Neuregulin signal, acting through ErbB4 receptors to drive blastema growth in *Hofstenia*. Neuregulin and/or EGFR signaling regulate injury-induced proliferative phenomena in many regenerative contexts and in diverse organisms (Ricci and Srivastava, 2018), including in planarians (Lei et al., 2016), zebrafish (Gemberling et al., 2015) (Rojas-Munoz et al., 2009), and axolotls (Farkas et al., 2016; Jensen et al., 2021) (Fig. 5C). Our findings confirm a likely ancient and widespread use of this pathway for promoting regeneration through the regulation of progenitor behavior.

## Materials and Methods

### Experimental Model

*Hofstenia miamia* were reared in artificial sea water prepared from Instant Ocean salt at an osmolarity of 37 ppt and pH 8.0 at 21°C. Adult animals were fed freshly hatched Artemia twice per week, and their embryos collected and grown in plastic petri dishes until hatching. Juvenile animals were used for all of the experiments, and these were grown with two changes to their water each week and fed with *Brachionus plicatilis* (L-type) marine Rotifers (Reed Mariculture) twice per week.

### RNA interference (RNAi)

Double-stranded RNA (dsRNA) was prepared through in vitro transcription as described previously (Clark and Petersen, 2023). dsRNA (1000 ng/μL) was microinjected on three consecutive days into the posterior gut region of animals using a Drummond microinjector, followed by amputation 3 hours later. Double-RNAi conditions involved mixing dsRNAs at equal concentrations to achieve a concentration of 1000 ng/μL. dsRNA targeting a gene sequence not present in the *Hofstenia* genome (*C. elegans unc-22*) was used as a negative control.

### Fluorescence In situ hybridization (FISH)

Animals were fixed with 4% paraformaldehyde in PBST (phosphate buffered saline with 0.1% Triton-X) for 1 hour at room temperature, washing with PBST, dehydration in methanol, and bleaching with 6% hydrogen peroxide. Fixed animals were stored at -20° until staining, then rehydrated, incubated in PBST, permeabilized with Proteinase K, refixed in 4% paraformaldehyde, followed by prehybridization hybridization buffer (50% formamide, 5xSSC, 1 ug/ml yeast RNA, 0.5% Tween) for 2 hours at 56°C, incubated 16 hours at 56°C in riboprobe mixed into hybridization buffer (50% formamide, 5xSSC, 1 ug/ml yeast RNA, 0.5% Tween, and 5% dextran sulfate), followed by washes in 2xSSC/0.1% Triton-X and 0.2xSSC/0.1% Triton-X. Samples were blocked with PBST with 10% Horse Serum, followed by incubation with anti-digoxigenin-HRP (1:1500) or anti-fluorescein-HRP antibody (1:1500) overnight at room temperature. Signal was developed using Tyramide Signal Amplification using homemade rhodamine-tyramide at 1:2000, and HRP inactivated between double-FISH steps using 100mM sodium azide as described (King and Newmark, 2013). Nuclei were stained were stained using 1:1000 Hoechst. Riboprobes were prepared by in vitro transcription from PCR-amplified templates containing T7 sequences, followed by transcription with T7 RNA polymerase and a nucleotide mixture containing digoxigenin-labeled or fluorescein-labeled UTP (Roche), resuspended in formamide and examined by agarose gel electrophoresis.

### Immunostaining

Animals were fixed and bleached as above, followed by blocking in PBST with 10% Horse Serum and incubation with rabbit anti-phospho-ser10-Histone3 (Cell Signaling Technology #3377S) at 1:500, then goat anti-rabbit-HRP (1:500) and tyramide signal development as above.

### Image Acquisition

Live animals were imaged with a Leica M210F dissecting microscope with a Leica DFC295 camera. Stained animals were imaged with a Leica Stellaris confocal microscope. FISH and immunostaining images show maximum projections from z-stacks. Adjustments to brightness and contrast were made using ImageJ or Adobe Photoshop.

### Cloning

Gene sequence identification was determined using the Hofstenia transcriptome and genome sequences(Gehrke et al., 2019; Srivastava et al., 2014). Primers for dsRNA and riboprobes were: *erbB4-1*(5’-GCAGCTCCAAAGAAACCAGT, 5’-AGTCGATCCCTCCAACGTTT-3’), *erbB4-2* (5’-GTGCACAGGGCCTAAAGAAA-3’, 5’-CCAAAGTGCAAATTGGAGGT-3’), *egf* 5’-TGCGATCCTCGAGAAAAACT-3’, 5’-TTGCTGTCAACTGCAAAACC-3’), *egr* (5’-GTTCCTCTCGCATAGCGAAT-3’, 5’-GTGGTTGGCGTCATACAATG-3’), *nrg-1* (5’-AGGTTAGACGGATCACTTCACT-3’, 5’-GCTACGCAACACAATTCATCA-3’), *nrg-2* (5’-CCTATGCAGCCGATGTTGTT-3’, 5’-CGATTTACCCAGCTCGCTTT-3’), *piwi-1* (5’-CCACCAAACATTCCGTTTTT-3’, 5’-GACCATCGGCAATTTCTTGT-3’), *sfrp1* (5’-TGCCGGAATTTCCTTAATTG-3’, 5’-GTGACCTACAGCCCCTTTGA), *tropomyosin* (5’-TTGTGGGGAGTGCCATTATT-3’,5’-TGAAACAAATGTTGGGATGAA-3’). A mixed stage library of total RNA was harvested from regenerating *Hofstenia* adults using Trizol and by reverse-transcription and PCR from a mixed-stage pool of *Hofstenia* total RNA. PCR products were TA-cloned into pGem-T-easy for generating templates for dsRNA or riboprobe synthesis.

### Quantification and Statistical Analysis

Counting of double-positive FISH cells was conducted by imaging through whole-animal z-stacks, and manually identifying *nrg-1+* cells, followed by assignment of the cell as muscle identity based on the presence of *tropomyosin* FISH signal detected surrounding the same Hoechst+ nucleus. Automated H3P counting and animal area analyses was performed in Fiji by an automated implementation of the Analyze Particles function. One-way ANOVA by Kruskal-Wallis test and

Dunnett’s tests were used in comparison of means of H3P+ cell density (number H3P+ cells per square millimeter) across multiple populations (R Studio).

### Protein structure prediction

Alphafold3 was used with default settings (https://alphafoldserver.com/) to predict structures of amino acids 28-81 of *Hofstenia* NRG-1 protein UEC49168.1 alone or also in complex with amino acids 25-868 of ERBB4-2 (predicted from contig “98056230|erbb4-2” contig in a previously reported *Hofstenia* transcriptome (Srivastava et al., 2014)) corresponding to the extracellular region. Quality scores were ipTM=0.85, pTM=0.61 for the ERBB4-2/NRG1 complex, and structures depicted are color-coded according to the predicted local distance difference test (pLDDT).

## Author contributions

BS: Investigation, Validation, Visualization, Roles/Writing - original draft, Writing - review & editing

RP: Investigation

HH: Investigation

CPP: Conceptualization, Funding acquisition, Project administration, Supervision, Roles/Writing - original draft, Writing - review & editing

## Acknowledgements

We thank Dr. Mansi Srivastava and the Srivastava lab members for the culture of *Hofstenia miamia* as well as extensive and generous advice on setup, experimental protocols, and feedback for work on this organism. We thank members of the Petersen lab for critical comments on the manuscript. This work was supported by National Institutes of Health grant NIGMS R35GM149280 (to C.P.P.). The funders had no role in study design, data collection and analysis, decision to publish, or preparation of the manuscript.

## Data and materials availability

All data are available in the main text or the supplementary materials.

---

**Figure S1.**
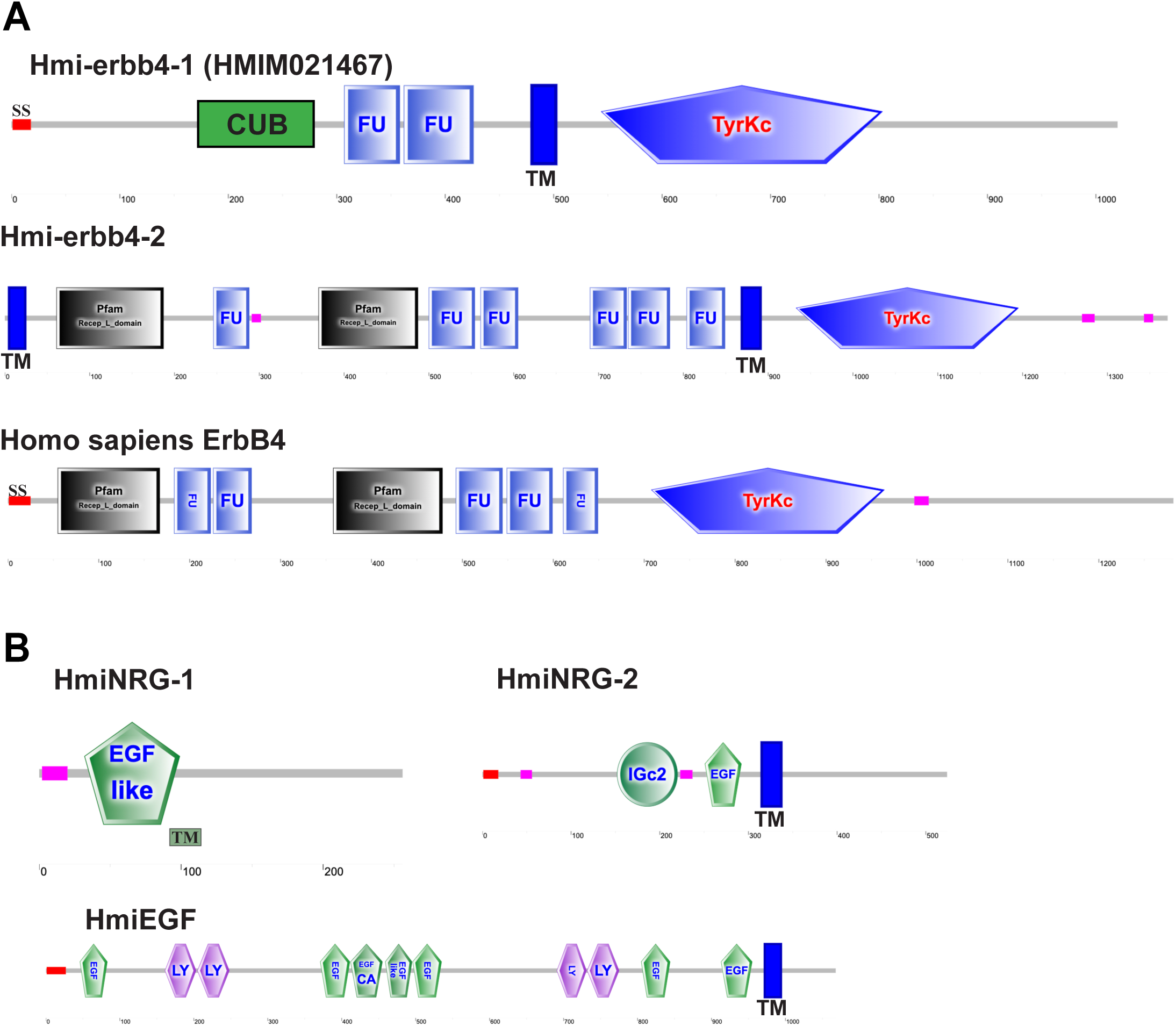
Domain structures of *Hofstenia* candidate EGFR and EGF-family genes. **(A)** *Hofstenia* ERBB4-1 has a well identifiable tyrosine kinase domain blasting to EGFRs, but its extracellular domain is divergent, possessing Furin domains but lacking any canonical Receptor L domains. Instead, the N terminal region of the protein may contain a highly diverged CUB domain (<u>C</u>omplement C1r/C1s, <u>U</u>egf, <u>B</u>mp1) identified below significance by Pfam (eval=27.7) but more confidently through a structural similarity search using Foldseek (https://search.foldseek.com/search), which recovered matches to the CUB domains of multiple secreted proteins (eg matches to Ovochymase-2 CUB domain, eval=7.96E-5; Match to Fibrocystin CUB domain, eval=2.54E-6). By contrast, ERBB4-2 has an overall canonical EGFR domain structure with Receptor L domains and Furin domains. The N-terminus lacks a signal sequence and instead contains a predicted N-terminal transmembrane domain (blue bars), suggesting this EGFR matures as a type II membrane protein with an N-terminal membrane anchor. (B) Domain structures of *Hofstenia* EGF family factors NRG-1, NRG-2, and EGF. NRG-1 contains a TM domain adjacent to the EGF-like domain. (A-B) Domains predicted by Pfam/Smart (http://smart.embl-heidelberg.de/).

**Figure S2.**
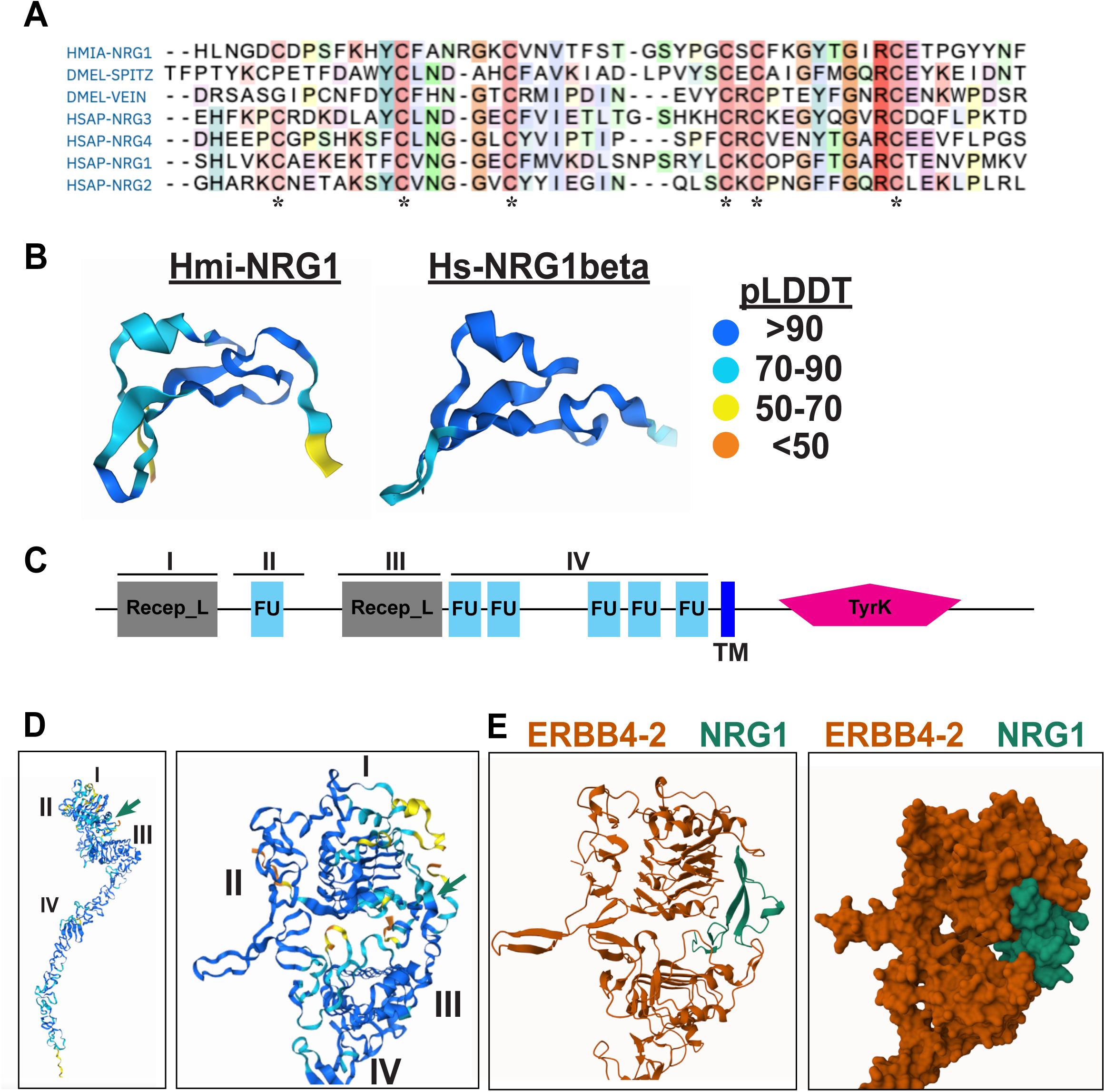
Prediction of interaction between NRG-1 and ERBB4-2. (A) Multiple sequence alignment (Clustal Omega) finds conservation of the canonical 6 cysteins present in *Hofstenia* NRG-1 (HMIA, *Hofstenia miamia*; DMEL *Drosophila melanogaster*, HSAP *Homo sapiens*). (B) Alphafold3 (https://alphafoldserver.com/) predictions of *Hofstenia* NRG-1 protein resemble the overall fold predicted for human NRG1beta. Color code displays predicted local distance difference test (pLDDT), with scores of >90 predicting highest accuracy (backbone and side chains) and scores of 70-90 indicating likely correct backbone with some sidechains misplaced. (C-D) Alphafold3 predicts interactions between the ERBB4-2 ectodomain and NRG-1. (C) The combined structure was used to annotate canonical EGFR domain regions (I-IV). Note ERBB4-2 has a somewhat extended region IV compared to other known EGFRs. (D) Left panel, Alphafold3 output of the combined structure. Right panel, zoom showing NRG-1 binding (green arrow) to a pocket composed of ERBB4-2 domains I and III. Colorscale depicts a strong overall pLDDT confidence (scaled as in panel B). (E) Ribbon (left) and space-filling (right) diagrams of the binding region, with ERBB4-2 (red) and NRG-1(green) colors indicated separately.

**Figure S3.**
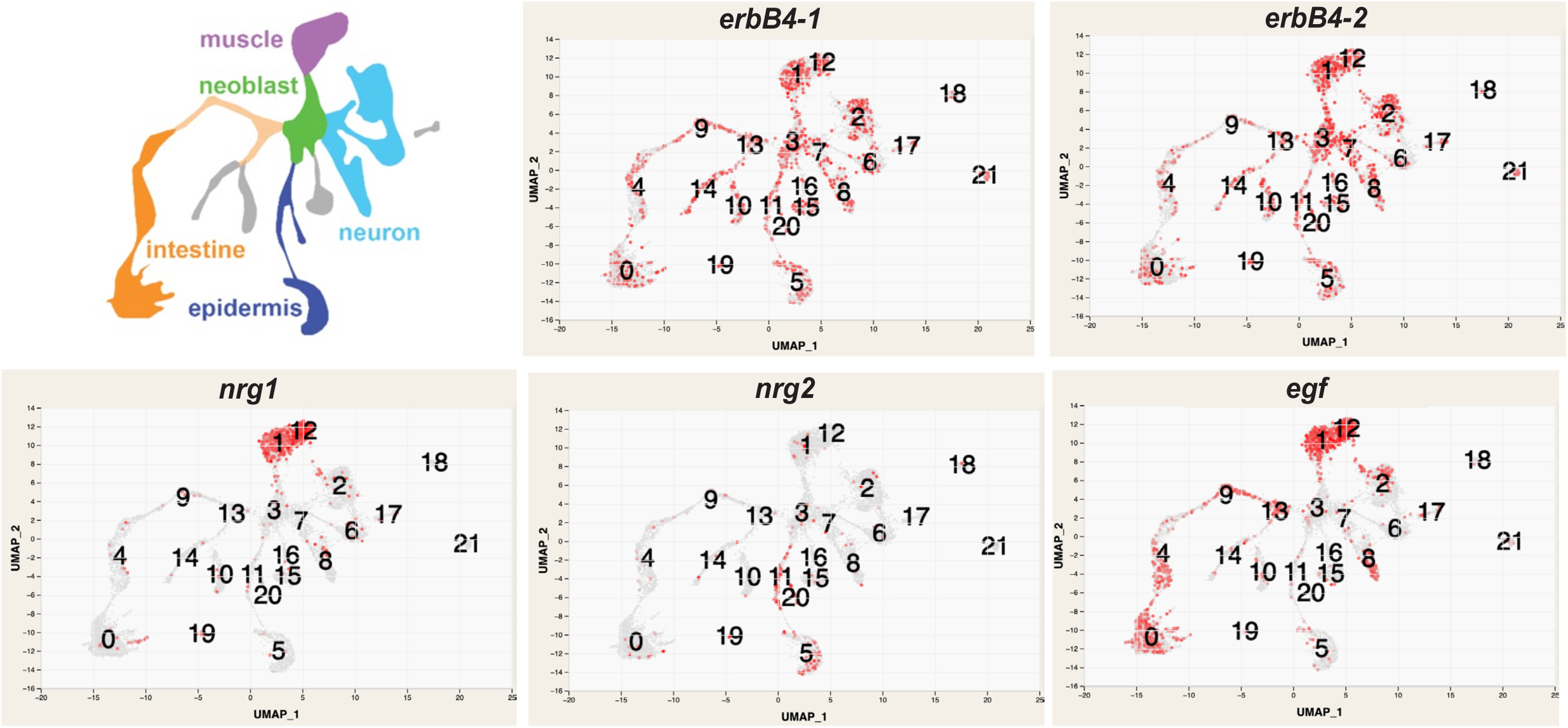
Expression of genes in this study within the *Hofstenia* single-cell atlas. Upper left, cartoon depicting cell types assigned to different categories. UMAP plots (https://dredge.ptgolden.org/sc/) show expression of *erbB4-1*, *erbB4-2*, *nrg1*, *nrg2*, and *egf* in a prior *Hofstenia* cell atlas (Hulett et al., 2023).

## References

Akheralie, Z., Scidmore, T.J., Pearson, B.J., 2023. aristaless-like homeobox-3 is wound induced and promotes a low-Wnt environment required for planarian head regeneration. Development 150.

Amiel, A.R., Rottinger, E., 2021. Experimental Tools to Study Regeneration in the Sea Anemone Nematostella vectensis. Methods Mol Biol 2219, 69–80.

Barberan, S., Cebria, F., 2019. The role of the EGFR signaling pathway in stem cell differentiation during planarian regeneration and homeostasis. Semin Cell Dev Biol 87, 45–57.

Barberan, S., Fraguas, S., Cebria, F., 2016. The EGFR signaling pathway controls gut progenitor differentiation during planarian regeneration and homeostasis. Development 143, 2089–2102.

Bork, P., Beckmann, G., 1993. The Cub Domain - a Widespread Module in Developmentally-Regulated Proteins. Journal of Molecular Biology 231, 539–545.

Bradshaw, B., Thompson, K., Frank, U., 2015. Distinct mechanisms underlie oral vs aboral regeneration in the cnidarian Hydractinia echinata. eLife 4, e05506.

Breen, C., Srivastava, M., 2024. A spreading, multi-tissue wound signal initiates whole-body regeneration. BioRxiv.

Breen, C.S., M., 2024. A spreading, multi-tissue wound signal initiates whole-body regeneration. BioRxiv.

Cannon, J.T., Vellutini, B.C., Smith, J., 3rd, Ronquist, F., Jondelius, U., Hejnol, A., 2016. Xenacoelomorpha is the sister group to Nephrozoa. Nature 530, 89–93.

Chan, A., Ma, S., Pearson, B.J., Chan, D., 2021. Collagen IV differentially regulates planarian stem cell potency and lineage progression. Proceedings of the National Academy of Sciences of the United States of America 118.

Citri, A., Yarden, Y., 2006. EGF-ERBB signalling: towards the systems level. Nat Rev Mol Cell Biol 7, 505–516.

Clark, E.G., Petersen, C.P., 2023. BMP suppresses WNT to integrate patterning of orthogonal body axes in adult planarians. PLoS genetics 19, e1010608.

Cote, L.E., Simental, E., Reddien, P.W., 2019. Muscle functions as a connective tissue and source of extracellular matrix in planarians. Nature communications 10, 1592.

Emili, E., Esteve Pallares, M., Romero, R., Cebria, F., 2019. Smed-egfr-4 is required for planarian eye regeneration. The International journal of developmental biology 63, 9–15.

Fan, Y., Chai, C., Li, P., Zou, X., Ferrell, J.E., Jr., Wang, B., 2023. Ultrafast distant wound response is essential for whole-body regeneration. Cell 186, 3606–3618 e3616.

Farkas, J.E., Freitas, P.D., Bryant, D.M., Whited, J.L., Monaghan, J.R., 2016. Neuregulin-1 signaling is essential for nerve-dependent axolotl limb regeneration. Development 143, 2724–2731.

Ferguson, K.M., 2008. Structure-based view of epidermal growth factor receptor regulation. Annu Rev Biophys 37, 353–373.

Fraguas, S., Barberan, S., Cebria, F., 2011. EGFR signaling regulates cell proliferation, differentiation and morphogenesis during planarian regeneration and homeostasis. Developmental biology 354, 87–101.

Gavino, M.A., Wenemoser, D., Wang, I.E., Reddien, P.W., 2013. Tissue absence initiates regeneration through follistatin-mediated inhibition of activin signaling. eLife 2, e00247.

Gehrke, A.R., Neverett, E., Luo, Y.J., Brandt, A., Ricci, L., Hulett, R.E., Gompers, A., Ruby, J.G., Rokhsar, D.S., Reddien, P.W., Srivastava, M., 2019. Acoel genome reveals the regulatory landscape of whole-body regeneration. Science 363.

Gemberling, M., Karra, R., Dickson, A.L., Poss, K.D., 2015. Nrg1 is an injury-induced cardiomyocyte mitogen for the endogenous heart regeneration program in zebrafish. eLife 4.

Hejnol, A., Obst, M., Stamatakis, A., Ott, M., Rouse, G.W., Edgecombe, G.D., Martinez, P., Baguna, J., Bailly, X., Jondelius, U., Wiens, M., Muller, W.E., Seaver, E., Wheeler, W.C., Martindale, M.Q., Giribet, G., Dunn, C.W., 2009. Assessing the root of bilaterian animals with scalable phylogenomic methods. Proc Biol Sci 276, 4261–4270.

Hulett, R.E., Kimura, J.O., Bolanos, D.M., Luo, Y.J., Rivera-Lopez, C., Ricci, L., Srivastava, M., 2023. Acoel single-cell atlas reveals expression dynamics and heterogeneity of adult pluripotent stem cells. Nature communications 14, 2612.

Hulett, R.E., Potter, D., Srivastava, M., 2020. Neural architecture and regeneration in the acoel Hofstenia miamia. Proc Biol Sci 287, 20201198.

Hulett, R.E., Rivera-Lopez, C., Gehrke, A.R., Gompers, A., Srivastava, M., 2024. A wound-induced differentiation trajectory for neurons. Proceedings of the National Academy of Sciences of the United States of America 121, e2322864121.

Jensen, T.B., Giunta, P., Schultz, N.G., Griffiths, J.M., Duerr, T.J., Kyeremateng, Y., Wong, H., Adesina, A., Monaghan, J.R., 2021. Lung injury in axolotl salamanders induces an organ-wide proliferation response. Developmental dynamics : an official publication of the American Association of Anatomists 250, 866–879.

Johnson, K., Bateman, J., DiTommaso, T., Wong, A.Y., Whited, J.L., 2018. Systemic cell cycle activation is induced following complex tissue injury in axolotl. Developmental biology 433, 461–472.

Kimura, J.O., Bolanos, D.M., Ricci, L., Srivastava, M., 2022. Embryonic origins of adult pluripotent stem cells. Cell 185, 4756–4769 e4713.

Kimura, J.O., Ricci, L., Srivastava, M., 2021. Embryonic development in the acoel Hofstenia miamia. Development 148.

King, R.S., Newmark, P.A., 2013. In situ hybridization protocol for enhanced detection of gene expression in the planarian Schmidtea mediterranea. BMC developmental biology 13, 8.

Lei, K., Thi-Kim Vu, H., Mohan, R.D., McKinney, S.A., Seidel, C.W., Alexander, R., Gotting, K., Workman, J.L., Sanchez Alvarado, A., 2016. Egf Signaling Directs Neoblast Repopulation by Regulating Asymmetric Cell Division in Planarians. Developmental cell 38, 413–429.

Lemke, G.E., Brockes, J.P., 1984. Identification and purification of glial growth factor. The Journal of neuroscience : the official journal of the Society for Neuroscience 4, 75–83.

Lindsay-Mosher, N., Chan, A., Pearson, B.J., 2020. Planarian EGF repeat-containing genes megf6 and hemicentin are required to restrict the stem cell compartment. PLoS genetics 16, e1008613.

Nanes Sarfati, D., Xue, Y., Song, E.S., Byrne, A., Le, D., Darmanis, S., Quake, S.R., Burlacot, A., Sikes, J., Wang, B., 2024. Coordinated wound responses in a regenerative animal-algal holobiont. Nature communications 15, 4032.

Newmark, P.A., Sanchez Alvarado, A., 2000. Bromodeoxyuridine specifically labels the regenerative stem cells of planarians. Developmental biology 220, 142–153.

Petersen, C.P., Reddien, P.W., 2009. A wound-induced Wnt expression program controls planarian regeneration polarity. Proceedings of the National Academy of Sciences of the United States of America 106, 17061–17066.

Petersen, C.P., Reddien, P.W., 2011. Polarized notum activation at wounds inhibits Wnt function to promote planarian head regeneration. Science 332, 852–855.

Philippe, H., Brinkmann, H., Copley, R.R., Moroz, L.L., Nakano, H., Poustka, A.J., Wallberg, A., Peterson, K.J., Telford, M.J., 2011. Acoelomorph flatworms are deuterostomes related to Xenoturbella. Nature 470, 255–258.

Plowman, G.D., Culouscou, J.M., Whitney, G.S., Green, J.M., Carlton, G.W., Foy, L., Neubauer, M.G., Shoyab, M., 1993. Ligand-specific activation of HER4/p180erbB4, a fourth member of the epidermal growth factor receptor family. Proceedings of the National Academy of Sciences of the United States of America 90, 1746–1750.

Poss, K.D., Shen, J., Nechiporuk, A., McMahon, G., Thisse, B., Thisse, C., Keating, M.T., 2000. Roles for Fgf signaling during zebrafish fin regeneration. Developmental biology 222, 347–358.

Poss, K.D., Tanaka, E.M., 2024. Hallmarks of regeneration. Cell stem cell 31, 1244–1261.

Poulet, A., Kratkiewicz, A.J., Li, D., van Wolfswinkel, J.C., 2023. Chromatin analysis of adult pluripotent stem cells reveals a unique stemness maintenance strategy. Sci Adv 9, eadh4887.

Ramirez, A.N., Loubet-Senear, K., Srivastava, M., 2020. A Regulatory Program for Initiation of Wnt Signaling during Posterior Regeneration. Cell Rep 32, 108098.

Raz, A.A., Srivastava, M., Salvamoser, R., Reddien, P.W., 2017. Acoel regeneration mechanisms indicate an ancient role for muscle in regenerative patterning. Nature communications 8, 1260.

Ricci, L., Srivastava, M., 2018. Wound-induced cell proliferation during animal regeneration. Wiley interdisciplinary reviews. Developmental biology 7, e321.

Ricci, L., Srivastava, M., 2021. Transgenesis in the acoel worm Hofstenia miamia. Developmental cell 56, 3160–3170 e3164.

Rink, J.C., Vu, H.T., Sanchez Alvarado, A., 2011. The maintenance and regeneration of the planarian excretory system are regulated by EGFR signaling. Development 138, 3769–3780.

Rio, C., Rieff, H.I., Qi, P., Khurana, T.S., Corfas, G., 1997. Neuregulin and erbB receptors play a critical role in neuronal migration. Neuron 19, 39–50.

Roberts-Galbraith, R.H., Newmark, P.A., 2013. Follistatin antagonizes activin signaling and acts with notum to direct planarian head regeneration. Proceedings of the National Academy of Sciences of the United States of America 110, 1363–1368.

Rojas-Munoz, A., Rajadhyksha, S., Gilmour, D., van Bebber, F., Antos, C., Rodriguez Esteban, C., Nusslein-Volhard, C., Izpisua Belmonte, J.C., 2009. ErbB2 and ErbB3 regulate amputation-induced proliferation and migration during vertebrate regeneration. Developmental biology 327, 177–190.

Ruiz-Trillo, I., Riutort, M., Littlewood, D.T., Herniou, E.A., Baguna, J., 1999. Acoel flatworms: earliest extant bilaterian Metazoans, not members of Platyhelminthes. Science 283, 1919–1923.

Scimone, M.L., Cote, L.E., Reddien, P.W., 2017. Orthogonal muscle fibres have different instructive roles in planarian regeneration. Nature 551, 623–628.

Scimone, M.L., Lapan, S.W., Reddien, P.W., 2014. A forkhead transcription factor is wound-induced at the planarian midline and required for anterior pole regeneration. PLoS genetics 10, e1003999.

Scimone, M.L., Wurtzel, O., Malecek, K., Fincher, C.T., Oderberg, I.M., Kravarik, K.M., Reddien, P.W., 2018. foxF-1 Controls Specification of Non-body Wall Muscle and Phagocytic Cells in Planarians. Current biology : CB 28, 3787–3801 e3786.

Srivastava, M., 2022. Studying development, regeneration, stem cells, and more in the acoel Hofstenia miamia. Current topics in developmental biology 147, 153–172.

Srivastava, M., Mazza-Curll, K.L., van Wolfswinkel, J.C., Reddien, P.W., 2014. Whole-body acoel regeneration is controlled by Wnt and Bmp-Admp signaling. Current biology : CB 24, 1107–1113.

Tahara, N., Brush, M., Kawakami, Y., 2016. Cell migration during heart regeneration in zebrafish. Developmental dynamics : an official publication of the American Association of Anatomists 245, 774–787.

Tewari, A.G., Owen, J.H., Petersen, C.P., Wagner, D.E., Reddien, P.W., 2019. A small set of conserved genes, including sp5 and Hox, are activated by Wnt signaling in the posterior of planarians and acoels. PLoS genetics 15, e1008401.

Vasquez-Doorman, C., Petersen, C.P., 2014. zic-1 Expression in Planarian neoblasts after injury controls anterior pole regeneration. PLoS genetics 10, e1004452.

Vogg, M.C., Owlarn, S., Perez Rico, Y.A., Xie, J., Suzuki, Y., Gentile, L., Wu, W., Bartscherer, K., 2014. Stem cell-dependent formation of a functional anterior regeneration pole in planarians requires Zic and Forkhead transcription factors. Developmental biology 390, 136–148.

Wenemoser, D., Lapan, S.W., Wilkinson, A.W., Bell, G.W., Reddien, P.W., 2012. A molecular wound response program associated with regeneration initiation in planarians. Genes & development 26, 988–1002.

Wenemoser, D., Reddien, P.W., 2010. Planarian regeneration involves distinct stem cell responses to wounds and tissue absence. Developmental biology 344, 979–991.

Witchley, J.N., Mayer, M., Wagner, D.E., Owen, J.H., Reddien, P.W., 2013. Muscle cells provide instructions for planarian regeneration. Cell Rep 4, 633–641.

Wurtzel, O., Cote, L.E., Poirier, A., Satija, R., Regev, A., Reddien, P.W., 2015. A Generic and Cell-Type-Specific Wound Response Precedes Regeneration in Planarians. Developmental cell 35, 632–645.

